# Independent Recruitment of the Mediator Tail Module to the *HO* Promoter Suggests Mediator Core Limits Coactivator Recruitment in *Saccharomyces cerevisiae*

**DOI:** 10.1101/766006

**Authors:** Robert M. Yarrington, Yaxin Yu, Chao Yan, Lu Bai, David J. Stillman

## Abstract

Mediator is an essential, multisubunit complex that functions as a transcriptional coactivator in yeast and other eukaryotic organisms. Mediator has four conserved modules, Head, Middle, Tail, and Kinase, and has been implicated in nearly all aspects of gene regulation. The Tail module has been shown to recruit the Mediator complex to the enhancer or UAS regions of genes via interactions with transcription factors, and the Kinase domain facilitates the transition of Mediator from the UAS/enhancer to the preinitiation complex via protein phosphorylation. Here we analyze expression of the *Saccharomyces cerevisiae HO* gene using a *sin4* Mediator Tail mutation that separates the Tail module from the rest of the complex; the *sin4* mutation permits independent recruitment of the Tail module to promoters without the rest of Mediator. Significant increases in recruitment of the SWI/SNF and SAGA coactivators to the *HO* promoter UAS were observed in a *sin4* mutant, along with increased gene activation. These results are consistent with recent studies that have suggested the Kinase module functions negatively to inhibit activation by the Tail. However, we found that Kinase module mutations did not mimic the effect of a *sin4* mutation on *HO* expression. This suggests that at *HO* the core Mediator complex (Middle and Head modules) must play a role in limiting Tail binding to the promoter UAS and gene activation. We propose that the core Mediator complex helps modulate Mediator binding to the UAS regions of genes to limit coactivator recruitment and ensure proper regulation of gene transcription.

## INTRODUCTION

Mediator is a large multisubunit transcriptional coactivator complex that is conserved throughout eukaryotes. Mediator was first identified in budding yeast as a bridge between general transcription factors (GTFs) and RNA Polymerase II (RNAPII; Thompson *et al*. 1993; Kim *et al*. 1994; Koleske and Young 1994). Currently, Mediator has been implicated in nearly all facets of gene regulation: transcriptional initiation and elongation (reviewed in Malik and Roeder 2000; Poss *et al*. 2013; Soutourina 2018), chromatin architecture (Allen and Taatjes 2015; Hsieh *et al*. 2015), mRNA processing and export (Huang *et al*. 2012; Schneider *et al*. 2015), and transcriptional memory (Zhang *et al*. 2013a; D’urso *et al*. 2016). Together these activities allow Mediator to modulate activation and expression required for proper gene regulation.

In yeast, mediator contains 25 subunits that are organized into four conserved modules: Head, Middle, Tail, and Kinase. The Head and Middle domains are essential for viability and thus make up the ‘core’ Mediator (Jeronimo and Robert 2017). Both of these modules interact with RNAPII as well as with GTFs required for transcriptional initiation and elongation. The Tail module attaches to the core Mediator through the Med14 (Rgr1) scaffold subunit (Tsai *et al*. 2014) and interacts with sequence-specific transcription factors that recruit the Mediator complex to the UAS or enhancers of genes (Bhoite *et al*. 2001; Kagey *et al*. 2010; Jeronimo and Robert 2014). The tail module is required for efficient recruitment of Mediator and Pol II to promoters (Knoll *et al*. 2018). The Kinase module is freely dissociable and attaches to the Middle module of Mediator via an interaction with its Med13 subunit. As part of the Mediator complex, the Kinase/CDK8 module antagonizes the function of the Tail module (Van De Peppel *et al*. 2005; Jeronimo and Robert 2017) and facilitates the transition of Mediator from the UAS to the preinitiation complex (PIC; Jeronimo *et al*. 2016; Petrenko *et al*. 2016).

Distinct forms of Mediator have been identified with and without the dissociable Kinase module (Poss *et al*. 2013), and additional stable forms have been identified through mutations of specific Mediator subunits. Deletion of the *SIN4 (MED16)* gene or a C-terminal truncation of the Med14 scaffold subunit each results in a stable tail-less Mediator complex and an independent Tail subcomplex composed of Med2, Pgd1 (Med3) and Gal11 (Med15) (Li *et al*. 1995). Yeast cells lacking the Mediator tail complex are viable but have pronounced defects in regulating a large subset of genes (Li *et al*. 1995; Larsson *et al*. 2013). This is likely due to the inability to recruit core Mediator and the CDK8/Kinase module to the UAS of these genes (Jeronimo *et al*. 2016; Petrenko *et al*. 2016). Changes in global chromatin structure (Macatee *et al*. 1997) and increased long-distance transcription factor activation (Dobi and Winston 2007) have also been reported during separation of the Tail subcomplex via disruption of *SIN4*.

Disruption of *SIN4* has been studied extensively at the *HO* promoter (Stillman *et al*. 1994; Tabtiang and Herskowitz 1998; Yu *et al*. 2000; Li *et al*. 2005). *HO* expression is under substantial regulation (Stillman 2013) and typically requires sequential activation at two distinct upstream regulatory regions, URS1 and URS2 (Nasmyth 1985; Cosma *et al*. 1999; Bhoite *et al*. 2001). Sequence-specific transcription factors Swi5 and SBF (Swi4/6 complex) bind to URS1 and URS2, respectively, and recruit the SWI/SNF, SAGA, and Mediator coactivator complexes. Loss of any of these factors results in defective *HO* activation. Disruption of *SIN4*, however, suppresses the requirement for SBF binding to URS2, and allows robust *HO* expression even when the Gcn5 catalytic subunit of SAGA is mutated (Yu *et al*. 2000). Mediator is initially recruited to the *HO* promoter via an interaction between Swi5 and the Gal11 Tail subunit (Bhoite *et al*. 2001), and it is likely that disruption of *SIN4* results in failure to recruit the ‘core’ Mediator and/or the Kinase/CDK8 and/or tail-less ‘core’ Mediator to the URS regions.

In this study, we further investigated the effects of disrupting *SIN4* at the *HO* promoter, and we explored the mechanism behind the resulting suppression of key regulatory events. We found that disrupting *SIN4* resulted in significant increases in both transcription factor and coactivator binding at the *HO* promoter, and that these increases are mostly due to prolonged persistence of these factors at the promoter. In agreement with this result, we also observed elevated and persistent nucleosome eviction during *HO* promoter activation. These results likely explain the suppression of multiple *HO* promoter mutants by *sin4Δ*. Surprisingly, we were unable to reproduce these results by mutating the catalytic subunit of the Kinase module, ruling out the simple model that the observed effects were due to loss of antagonistic effects of the Kinase module on Tail function. Rather, we found that elevated coactivator binding and suppression of promoter mutants were completely dependent on the presence of the Mediator Tail subcomplex. As we do not observe these effects with whole Mediator, we propose that the core Mediator must restrict the binding of either the Tail subcomplex or other transcription factors/coactivators to limit promoter activation and ensure proper regulation of gene transcription.

## MATERIALS AND METHODS

All yeast strains used in this study are listed in Table S1 and are isogenic in the W303 background (Thomas and Rothstein 1989). Standard genetic methods were used for strain construction (Rothstein 1991; Sherman 1991). The 5X-sbf, +700, and +1300 *HO* promoter mutants were described in (Yarrington *et al*. 2015), and the *ho(m-2700)* mutation was described in (Yu *et al*. 2016). C-terminal epitope tags were added as described (Knop *et al*. 1999), using plasmids pZC03 (pFA6a-TEV-6xGly-V5-HIS3MX; Addgene plasmid #44073) and pZC13 (pFA6a-TEV-6xGly-V5-HphMX; Addgene plasmid #44085), provided by Zaily Connell and Tim Formosa, and plasmid pYM6 (Knop *et al*. 1999), provided by Elmar Schiebel. Strain YTT1722 with a *SWI2:FLAG(3):KanMX* tag (Kim *et al*. 2006) was provided by David Clark, and the Marker Swap method (Voth *et al*. 2003) was used to convert it to *SWI2:FLAG(3):Nat MX* using plasmid pAG25 (Goldstein and Mccusker 1999) provided by John McCusker. Oligos used in strain construction are available upon request.

Cell cycle synchronization was performed by galactose withdrawal and re-addition with a *GALp::CDC20* strain grown at 25°C in YP medium containing 2% galactose and 2% raffinose (Bhoite *et al*. 2001). A high degree of synchrony was confirmed by examination of budding indices and analysis of cycle-regulated mRNAs. In all other experiments, cells were grown at 30°C in YPAD medium (Sherman 1991).

ChIPs were performed as described (Bhoite *et al*. 2001; Voth *et al*. 2007) using mouse monoclonal antibody to the V5 epitope (SV5-Pk1, Abcam) or anti-histone H3 (07-690, Upstate), and antibody-coated magnetic beads (Rabbit and Pan Mouse IgG beads, Life Technologies). Samples prepared for ChIPs were cross-linked in 1% formaldehyde overnight on ice. ChIP assays were analyzed by real time qPCR as described (Eriksson *et al*. 2004). ChIP qPCR primers are available upon request. H3 samples were first normalized to the ChIP signal at the IGR-I gene-free reference region on chromosome I (Mason and Struhl 2005), while Swi4-V5 ChIP samples were first normalized to the *CLN1* promoter, and then both types of ChIPs were normalized to their respective input DNA sample. Unless otherwise noted, error bars reflect the standard deviation of at least three biological samples. P-values were calculated by paired t-tests.

RNA was isolated from either synchronized or logarithmically growing cells, and *HO* mRNA levels were measured by RT-qPCR as described (Voth *et al*. 2007). For all logarithmically grown strains, RNA expression was normalized to *RPR1* expression and graphed relative to wild-type expression. For the synchrony experiment, RNA expression was normalized to *RPR1* expression and graphed relative to the peak WT expression. Unless otherwise noted, error bars reflect the standard deviation of at least three biological samples. P-values were calculated by paired t-tests. RT-qPCR primers are available upon request.

Single cell analysis of *HO* expression was performed by time-lapse fluorescence microscopy as described (Zhang et al., 2013).

Data Availability and Reagent Sharing. Strains are listed in Table S1 and are available upon request. Four Supplemental Figures and one Supplemental Table was uploaded to the GSA Figshare portal.

## RESULTS

### Disruption of *SIN4* can rescue *HO* promoter mutants that affect activation of URS2

We have previously demonstrated that *sin4Δ* can rescue *HO* expression when the Swi6 subunit of SBF is also disrupted (Yu *et al*. 2000). The SBF complex, however, is an important transcriptional regulator with numerous binding sites at genes controlling the G1/S transition (Andrews and Herskowitz 1989), and our previous suppression result could have been impacted by alterations to the yeast cell cycle. To demonstrate the impact of *sin4Δ* specifically at the *HO* gene, we made use of *HO* promoter mutants that either eliminated SBF sites at the left-half of URS2 (URS2L) or increased the distance between URS1 and URS2 (Fig 1; Yarrington *et al*. 2015). Both of these mutation classes drastically reduce SBF binding to the entire URS2 region with corresponding decreases in *HO* expression (Yarrington *et al*. 2015).

**Fig 1.**
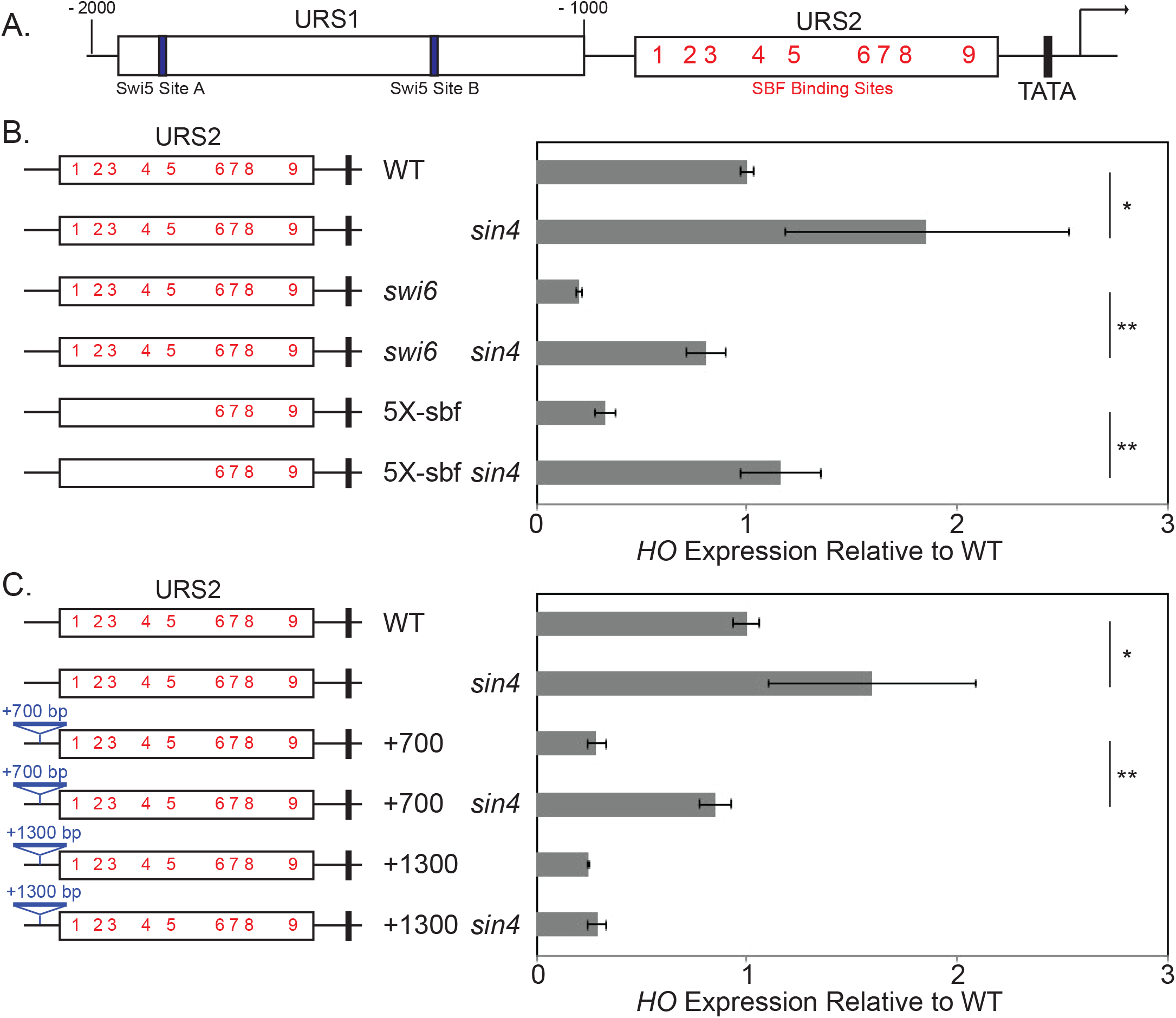
A *sin4* mutation rescues expression of *HO* promoter mutants. **A.** The diagram shows the structure of the *HO* promoter, including the two Swi5 binding sites at URS1 and the nine SBF binding sites in URS2. **B.** *HO* mRNA levels were measured for the various mutant *HO* promoters indicated on the left, in either wild type, *sin4, swi6*, or *swi6 sin4* mutants. The error bars reflect the standard deviation of three biological samples. \**p* < 0.05, \*\**p* < 0.01. **C.** *HO* mRNA levels were measured for the various mutant *HO* promoters indicated on the left, in either wild type or *sin4* mutants. The error bars reflect the standard deviation of three biological samples. \**p* < 0.05, \*\**p* < 0.01.

*HO* expression results confirmed previous findings that SBF is essential for WT-levels of *HO* expression and that *sin4Δ* is capable of suppressing a *SWI6* disruption (Fig 1B). Interestingly, we observed that *sin4Δ* alone resulted in elevated *HO* expression compared to WT. Additionally, mutation of the five SBF sites at the left-half of URS2 (the 5x-sbf construct, Fig 1B) or inserting 700bp of *CDC39* exon sequence between URS1 and URS2 (+700, Fig 1C) resulted in deceases in *HO* expression similar to that seen in a *swi6* mutant. In both cases, however, disruption of *SIN4* rescued *HO* expression to levels similar to that of WT. As mutating the *HO* promoter does not affect cell cycle progression, these results indicate that suppression of *swi6Δ* by *sin4Δ* is not an artifact of cell cycle impairment. Furthermore, as SBF binding to the right half of URS2 (URS2R) is essential for *HO* expression (Yarrington *et al*. 2015), these results suggest that *sin4Δ* or Mediator Tail separation allows SBF binding to URS2R under circumstances when it is normally prohibited, such as mutation of SBF sites at URS2L or insertion of 700bp between URS1 and URS2. This suppression is not without limits as the reduced *HO* expression caused by a 1300bp insertion between URS1 and URS2 (+1300) was not suppressed by *sin4Δ* (Fig 1C).

### Disruption of *SIN4* rescues SBF binding to URS2 in *HO* promoter mutants

We next wanted to test the hypothesis that disruption of *SIN4* rescues SBF binding to URS2 in our *HO* promoter mutants. We have previously shown that mutation of SBF sites at URS2L or increasing the distance between URS1 and URS2 inhibits binding of SBF to the entire URS2 region (Yarrington *et al*. 2015). To test whether *sin4Δ* can rescue SBF binding, we performed a Swi4-V5 ChIP in strains with these *HO* promoter mutations and probed SBF binding at both URS2L and URS2R (Fig 2).

**Fig 2.**
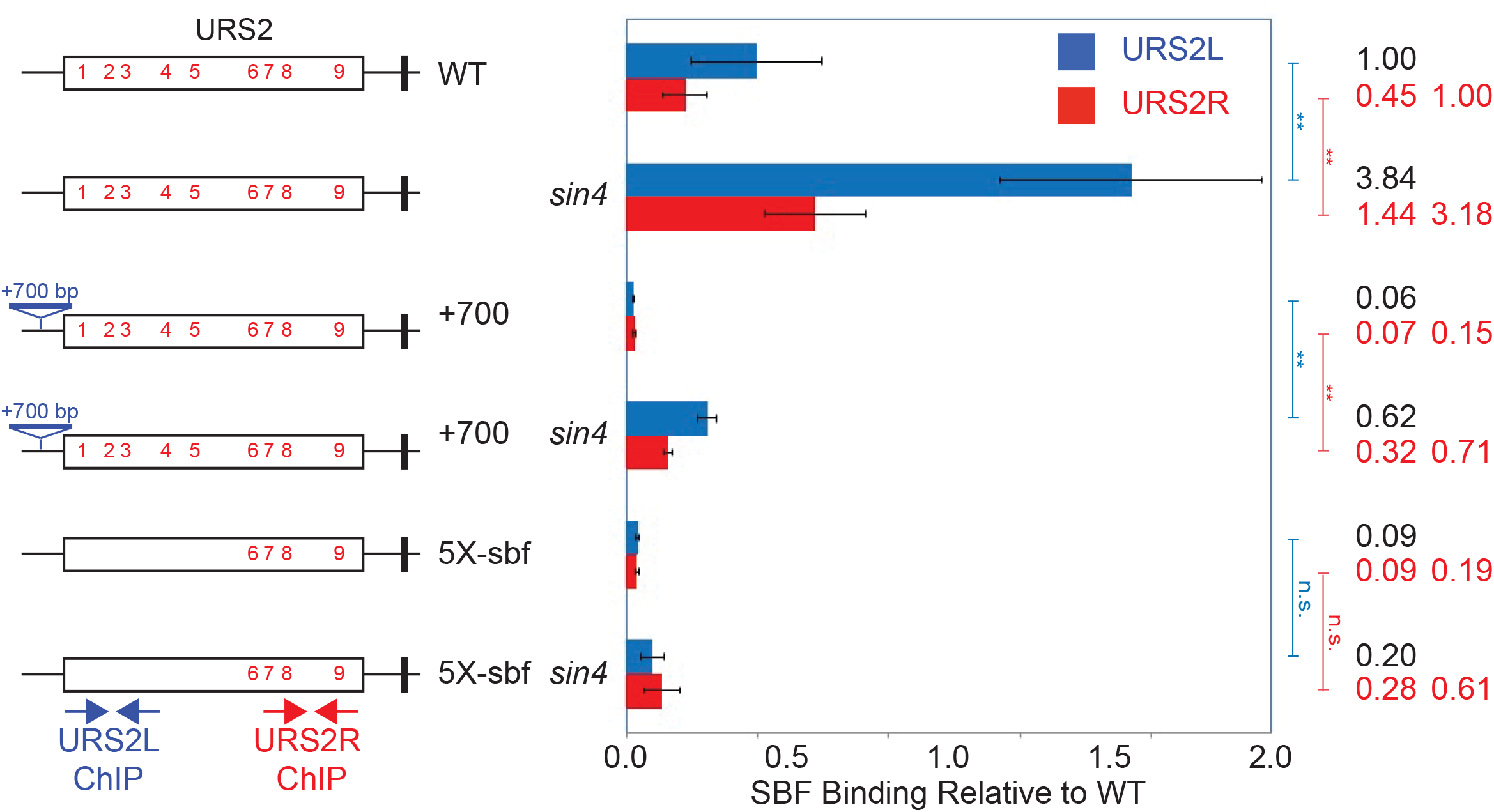
A *sin4* mutation rescues SBF binding in *HO* promoter mutations. SBF binding was measured by ChIP assays of the Swi4-V5 subunit of SBF to the various mutant *HO* promoters indicated on the left, in either wild type or *sin4* mutants. SBF binding was measured to the left (blue) or the right (red) parts of URS2, using the primers indicated on the diagram. The first column on the right displays enrichment values relative to the native URS2L part of the promoter, and the second column on the right displays enrichment values relative to the native URS2R part of the promoter. The error bars reflect the standard deviation of three biological samples. \*\**p* < 0.01.

As expected, SBF enrichment at URS2L is over 2-fold higher than at URS2R in WT cells, despite similar numbers of SBF sites (Fig 2). Preferential enrichment to URS2L has been observed previously (Takahata *et al*. 2011; Yarrington *et al*. 2015) and is due to closer proximity to remodeling events initiated at the upstream URS1 region. Interestingly, a *sin4Δ* mutation results in ~3-fold increase in SBF binding compared to WT, at both URS2L and URS2R, and this elevated SBF binding likely explains the similar increase in *HO* expression of this mutant. In the +700 and 5X-sbf *HO* promoter mutants, disruption of *SIN4* resulted in similar increases in SBF binding to URS2R to levels around two-thirds that of WT and approximately 4-fold higher than that of the single mutants without *sin4Δ* (Fig 2). In cell cycle synchrony experiments, we also see prolonged SBF binding to *HO* in the *sin4Δ* mutant, compared to wild type (Supplemental Fig S1). These results support the model that *sin4Δ* or separation of the Mediator Tail subcomplex enhances SBF binding to URS2 even when combined with mutations that normally block SBF binding.

### *sin4*Δ-mediated suppression of *HO* is dependent on Swi5 and Gal11

We have shown that separation of the Mediator Tail subcomplex enhances SBF binding to URS2 and suppresses defects in *HO* expression in certain promoter mutants, but the mechanism behind this suppression remains largely unknown. To investigate this mechanism further, we combined *sin4Δ* with other mutations known to affect *HO* expression.

We first focused on the approximately 2 to 3-fold increase in both *HO* expression and SBF binding observed in *sin4Δ* relative to WT. Similar increases occur when the gene for the daughter-specific inhibitor Ash1 factor is disrupted (Takahata *et al*. 2011; Stillman 2013). Combining *sin4Δ* with an *ASH1* disruption, however, revealed that these two mutations are additive for *HO* expression (Supplemental Fig S2) and are therefore not working together in the same pathway.

We next investigated the Swi5 transcription factor that recruits Mediator to URS1 to initiate *HO* activation (Bhoite *et al*. 2001). Combining *sin4Δ* with a *SWI5* disruption blocked promoter activation similarly to that of *swi5Δ* alone, and *swi5Δ sin4Δ* failed to activate expression from the 5X-sbf *HO* promoter mutant with five SBF site mutations (Fig 3A). This result indicates that *sin4Δ*-mediatied suppression does not bypass normal activation of the *HO* promoter. We further probed the relationship between *sin4Δ* and Swi5 by examining Swi5-V5 enrichment levels during logarithmic growth by ChIP at its two binding sites in URS1, site A and B (Tebb *et al*. 1993). Comparing Swi5-V5 enrichment in WT and *sin4Δ* revealed an ~50% increase in Swi5 binding at both sites when *SIN4* is disrupted (Fig 3B). Swi5 binding normally peaks 20 minutes after release from a G2/M arrest and quickly dissipates as the cell cycle progresses (Takahata et al., 2009). To determine whether the increase in Swi5-V5 enrichment in *sin4Δ* was due to greater peak binding at 20 minutes after release or to a change in binding kinetics, we synchronized cells using *GALp:CDC20* arrest and examined Swi5-V5 binding after release. Total Swi5 binding at 20 minutes post release was largely similar between WT and *sin4Δ* with the major difference being significant persistence of the Swi5 factor at URS1 in the *sin4* mutant (Fig 3C).

**Fig 3.**
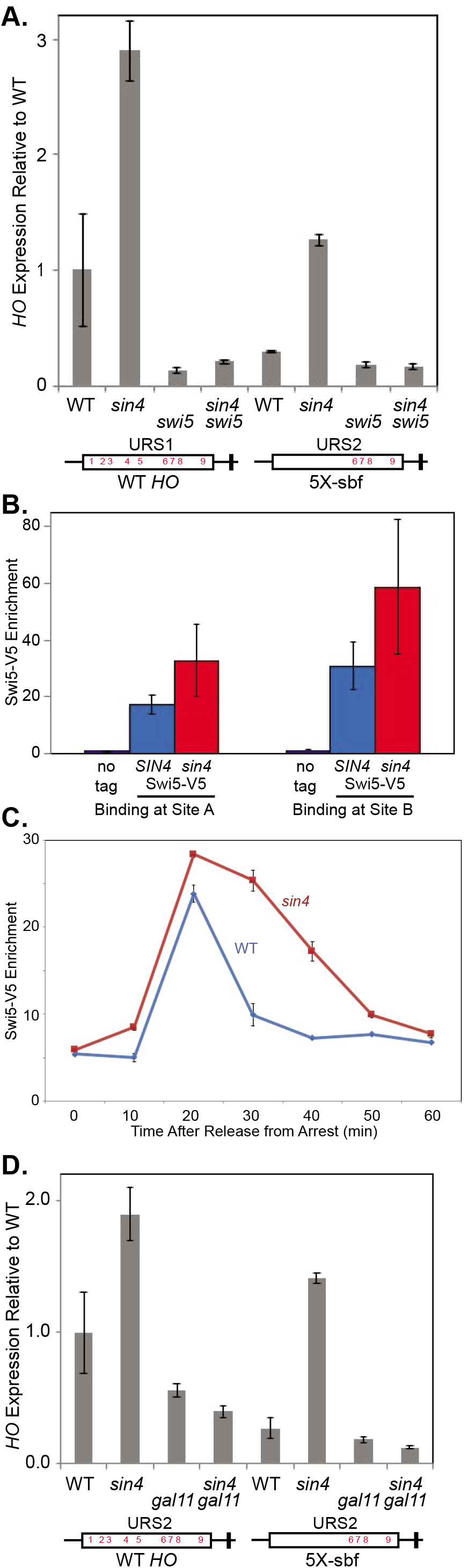
The *sin4* suppression of *HO* expression is dependent on Swi5 and Gal11. **A.** The *sin4* suppression of *HO* expression is dependent on Swi5. *HO* mRNA levels were measured for the wild type *HO* promoter (left four columns) or the 5X-sbf *HO* promoter mutant (right four columns), in either wild type, *sin4, swi5*, or *sin4 swi5* strains. The error bars reflect the standard deviation of two biological samples. **B.** Swi5-V5 binding by ChIP is higher in *sin4* mutants in log phase cells at both Site A (–1819) and Site B (–1308); see diagram in Fig 1A. The error bars reflect the standard deviation of three biological samples. **C.** Swi5-V5 binding by ChIP at Site B is higher in *sin4* mutants in cells synchronized with a *GALp:CDC20* arrest and release. The error bars reflect the standard deviation of PCR replicates. **D.** The *sin4* suppression of *HO* expression is dependent on Gal11. *HO* mRNA levels were measured for the wild type *HO* promoter (left four columns) or the 5X-sbf *HO* promoter mutant (right four columns), in either wild type, *sin4, gal11*, or *sin4 gal11* strains. The error bars reflect the standard deviation of two biological samples.

It has also been shown previously that Swi5 directly interacts with the Gal11 subunit of the Tail subcomplex, and that Mediator fails to bind when *GAL11* is disrupted (Bhoite *et al*. 2001). Disruption of *SIN4*, however, separates the Tail subcomplex from the rest of Mediator and it remains capable of binding independently of the rest of Mediator (Zhang *et al*. 2004; Ansari and Morse 2012). We performed a ChIP experiment to determine whether Tail and Core Mediator binding are also independent at *HO*. Srb4(Med17) is a subunit of the Head module of Mediator, and we have previously shown that Srb4 binds to both URS1 and URS2 of *HO* (Bhoite *et al*. 2001). Here we show that Srb4-V5 binding to *HO* is eliminated by a *sin4Δ* mutation (Supplemental Fig S3), consistent with separation of the Tail module from the rest of Mediator in a *sin4Δ* mutant. To determine whether the Tail subcomplex was required for *sin4Δ*-mediated suppression, we next examined *HO* expression when *SIN4* and *GAL11* were disrupted either independently or together. The double mutant did not have elevated *HO* expression in the WT promoter and failed to rescue expression of the 5x-sbf site mutant *HO* promoter, indicating that *sin4Δ*-mediated suppression is dependent on functional Gal11 (Fig 3D). Furthermore, disruptions of other Tail subunits, *PGD1* and *MED2*, also blocked *sin4Δ*-mediated suppression (Supplemental Fig S4). These results suggest that it is the independent recruitment of the Tail subcomplex, rather than the loss of core Mediator or its associated Kinase module, that is responsible for the observed suppression.

### *sin4Δ*-mediated suppression is not dependent on the Mediator CDK module

The current model for Mediator-mediated promoter regulation proposes that the Kinase module antagonizes the function of the Tail module (Van De Peppel *et al*. 2005; Jeronimo and Robert 2017). Disruption of *SIN4* leads to independent recruitment of the Tail subcomplex and loss of the Kinase/CDK8 module (Jeronimo *et al*. 2016; Petrenko *et al*. 2016), and our data is consistent with hyperactivation due to the loss of an inhibitor of promoter activation. Furthermore, the Srb10 catalytic subunit of the Kinase module has been shown to target Swi5 for degradation (Kishi *et al*. 2008), and failure to recruit the Kinase module might cause Swi5 protein levels to persist in a way congruent with our findings. This model, however, is not consistent with the requirement of a functional Tail subcomplex for suppression. To better understand the role of the Kinase module in *sin4Δ*-mediated hyperactivation of the *HO* promoter and suppression of promoter mutants, we examined *HO* expression when *SIN4* and *SRB10* were disrupted either independently or together. As expected, disruption of *SIN4* resulted in elevated expression of the native promoter and suppression of 5X-sbf promoter to WT levels (Fig 4). Loss of Srb10, however, failed to show an appreciable effect on either the native or the mutant promoter, and *sin4Δ srb10Δ* double mutants produced expression results only modestly different from those of *sin4Δ* single mutants (Fig 4). These results indicate that loss of Kinase module activity has a minimal impact on *HO* expression. Additionally, these results further support the interpretation that *sin4Δ*-mediated hyperactivation of the *HO* promoter is due to independent recruitment of the Tail subcomplex rather than loss of the Kinase module associated with Mediator.

**Fig 4.**
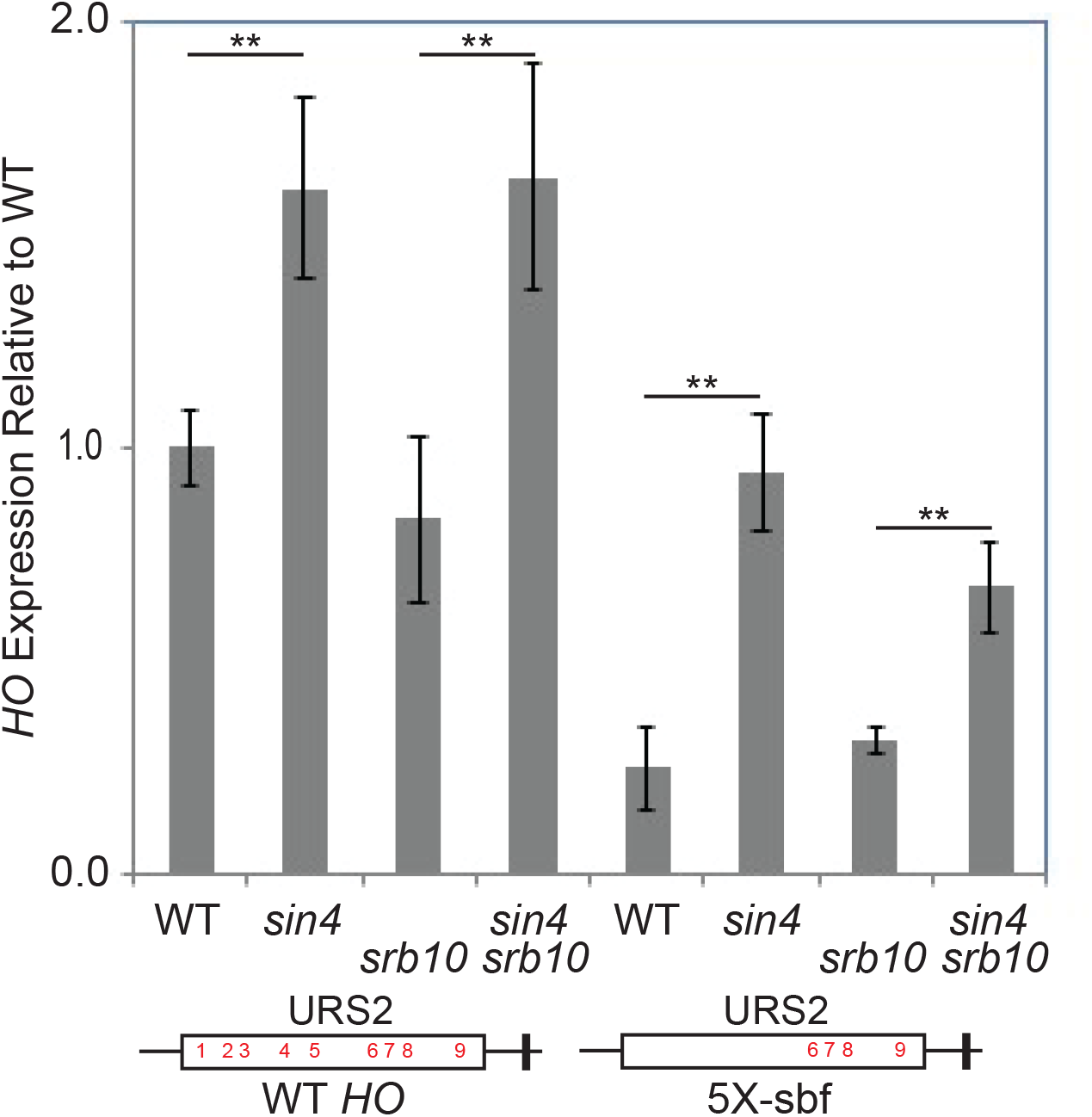
Loss of the Kinase module with an *srb10* mutation does not affect *HO* expression. *HO* mRNA levels were measured for the wild type *HO* promoter (left four columns) or the 5X-sbf *HO* promoter mutant (right four columns), in either wild type, *sin4, srb10*, or *sin4 srb10* strains. The error bars reflect the standard deviation of four biological samples. \*\**p* < 0.01.

### Disruption of *SIN4* results in elevated coactivator enrichment at the *HO* promoter

Swi5 recruits the SWI/SNF, SAGA, and Mediator coactivator complexes to URS1, and SBF recruits these same complexes to URS2 (Takahata *et al*. 2009). Disruption of *SIN4* causes increased enrichment of both of these DNA-binding factors and may similarly affect their ability to recruit coactivators to the *HO* promoter. Elevated and/or persistent recruitment of coactivators to the *HO* promoter could explain the hyperactivation observed with disrupting *SIN4*. To examine this possibility, we performed ChIP to measure recruitment of Swi2-V5 of the SWI/SNF complex, Gcn5-V5 of the SAGA complex, and Gal11-V5 of the Mediator complex. As Gal11 is part of the Mediator Tail module, in the *sin4Δ* mutant we were only examining recruitment of the Tail subcomplex.

We first examined binding of the coactivator complexes to *HO* URS1 during logarithmic growth and found that disrupting *SIN4* caused coactivator enrichment to increase for all three complexes, ranging from 1.6 to 2-fold enhanced enrichment (Fig 5A). A functional Tail module was required for the hyperactivation and suppression by *sin4Δ* (Fig 3D), and we next investigated whether this Tail-dependence held true for coactivator complex recruitment. To test this possibility, we examined Swi2-V5 binding when *SIN4* and *GAL11* were disrupted either independently or together. Coactivator binding is interdependent (Takahata *et al*. 2011), and we observed an expected small decrease in Swi2-V5 enrichment in the *gal11Δ* single mutant. In the *gal11Δ sin4Δ* double mutant, however, we saw complete loss of the near 4-fold enrichment observed with the *sin4Δ* single mutant (Fig 5B). These results suggest that independent recruitment of the Tail subcomplex is necessary and sufficient to increase recruitment of SWI/SNF to the *HO* promoter.

**Fig 5.**
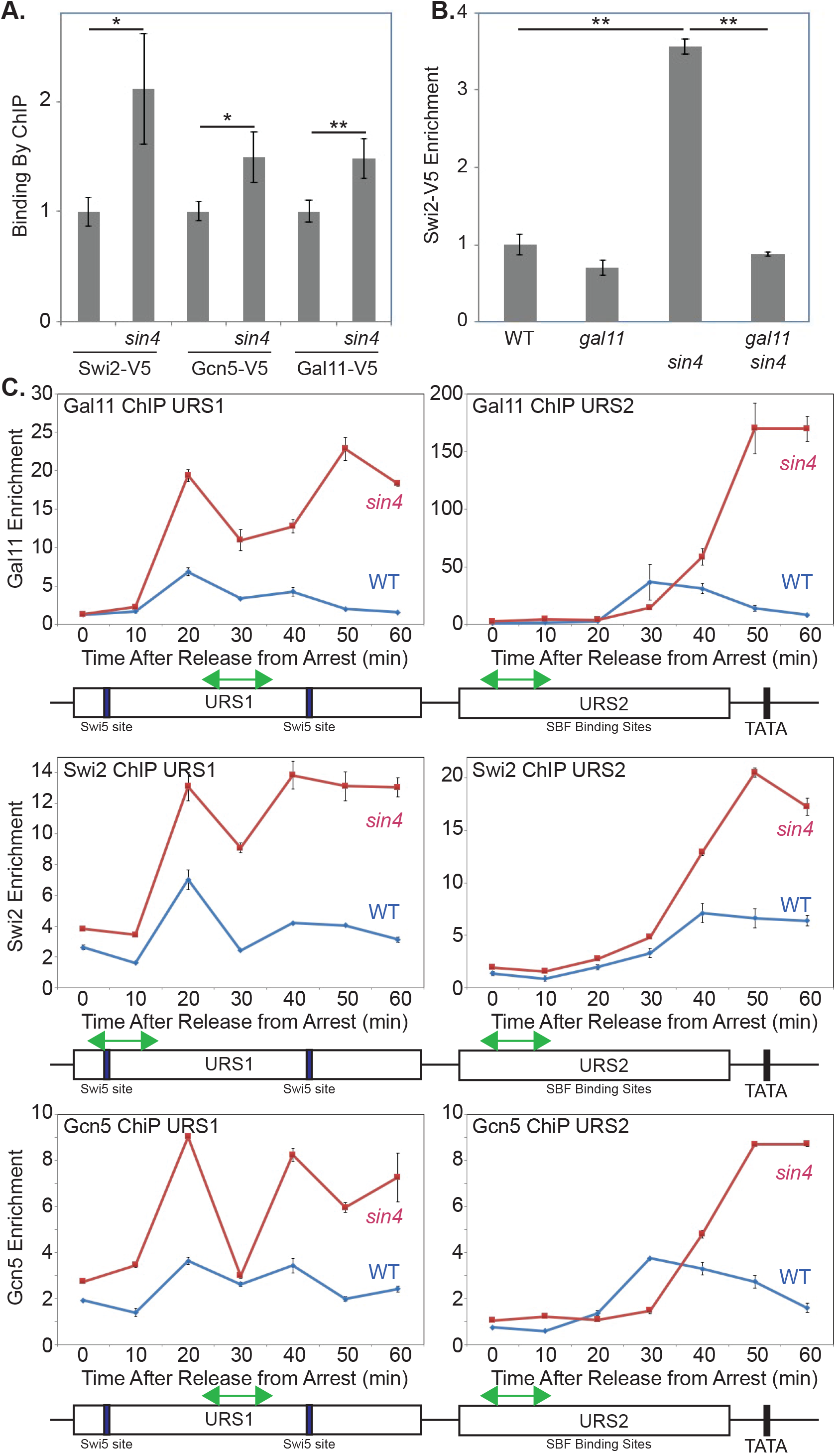
A *sin4* mutation increases coactivator binding to the *HO* promoter. **A.** The *sin4* mutation increases Swi2-V5, Gcn5-V5, and Gal11-V5 binding to the *HO* promoter as measured by ChIP. The error bars reflect the standard deviation of three biological samples. \**p* < 0.05, \*\**p* < 0.01. **B.** The increase in SWI/SNF binding measured by ChIP assays with Swi2-V5 are dependent upon Gal11. The error bars reflect the standard deviation of three biological samples. \*\**p* < 0.01. **C.** The increased coactivator binding in a *sin4* mutant persists during the cell cycle. Wild type (blue) and *sin4* (red) cells with a *GALp:CDC20* allele and either a Gal11-V5, a Swi2-V5, or a Gcn5-V5 epitope tag were synchronized by galactose withdrawal and readdition, and factor binding was measured by ChIP during the cell cycle. The ChIP signal is plotted as a function of time after release from the G2/M arrest. The left and right panels show binding at URS1 and URS2, respectively, with the positions of the PCR primers shown by green arrows. The error bars reflect the standard deviation of PCR replicates.

Additionally, the Swi5 factor persists at the *HO* promoter in a *SIN4* disruption (Fig 3C), and this persistence may affect the binding kinetics of recruited coactivators. To examine this possibility, we synchronized cells using *GALp:CDC20* arrest and release, and performed ChIP on the V5-tagged coactivator subunits. For all three coactivator complexes, we observed greater and prolonged enrichment to URS1 (Fig 5C). Interestingly, coactivator enrichment at URS1 continues well past Swi5 binding at URS1 (compare Fig 3C and 5C). We also found enhanced and persistent coactivator enrichment at URS2, and this recruitment appears to be in good agreement with altered SBF binding in *sin4Δ* (Supplemental Fig S1).

### Disruption of *SIN4* enhances nucleosome eviction at the *HO* promoter

*HO* experiences waves of nucleosome eviction along the length of its promoter during normal activation (Takahata *et al*. 2009). These waves of nucleosome eviction, however, are dependent on transient recruitment of transcription factors and coactivators whose binding kinetics are altered in a *SIN4* disruption. To determine whether *sin4Δ* affects nucleosome eviction at *HO*, we performed a H3 histone ChIP, examining H3 occupancy every 100 to 200 bp between URS1 and URS2R, after *GALp:CDC20* arrest and release.

Nucleosome eviction at URS1 was nearly identical between WT and *sin4Δ* for the first 20 minutes after release (Fig 6A). In agreement with altered coactivator recruitment (Fig 5C), however, nucleosome eviction appeared to continue beyond 20 minutes in *sin4Δ* leading to greater and more persistent eviction of nucleosomes. Typically, nucleosome occupancy at URS1 is mostly repopulated after 60 minutes post release (Takahata *et al*. 2009) but repopulation in *sin4Δ* was significantly delayed (Fig 6B, compare Wild Type and *sin4Δ* −1208 and −1109). Nucleosome eviction at URS2 was also affected, with much greater nucleosome depletion observed after 30 minutes post release in *sin4Δ* (Fig 6A and B). For example, the H3 occupancy data clearly shows that at 50 min the *sin4* mutant has marked delay in nucleosome repopulation along the *HO* promoter (Fig 6C). These results indicate that independent recruitment of the Tail subcomplex leads to enhanced nucleosome eviction at the *HO* promoter, presumably due to increased and persistent recruitment of the SWI/SNF complex.

**Fig 6.**
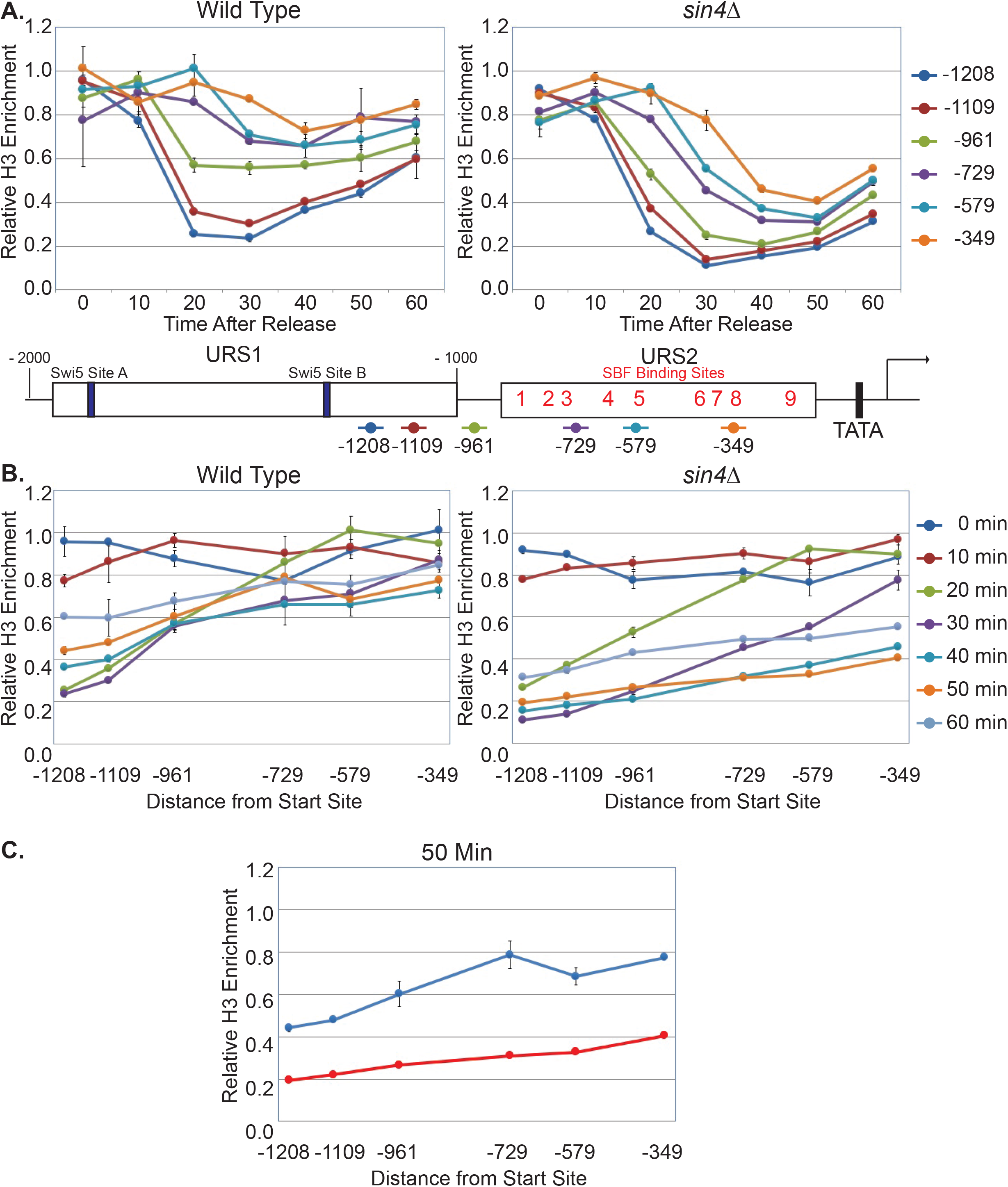
A *sin4* mutation prolongs nucleosome eviction at the *HO* promoter. Wild type (left) and *sin4* (right) cells with a *GALp:CDC20* arrest allele were synchronized by galactose withdrawal and re-addition, and H3 occupancy at the *HO* promoter was measured by ChIP during the cell cycle using primers spaced along the *HO* promoter. The error bars reflect the standard deviation of PCR replicates. **A.** The H3 ChIP is plotted as a function of time after release from the G2/M arrest, with the centers of the PCR intervals amplified listed on the right. The positions of the PCR amplicons are shown on the promoter map below. **B.** The data from panel A is plotted as a function of the distance along the promoter, with the time points listed on the right. **C.** The 50 min time points for wild type and *sin4* are plotted as a function of the distance along the promoter.

### Disruption of *SIN4* alters coactivator interdependence and increases the probability of *HO* promoter activation

Mutations in the catalytic subunits of SWI/SNF and SAGA greatly reduce *HO* expression and recruitment of other coactivators (Cosma *et al*. 1999; Mitra *et al*. 2006; Takahata *et al*. 2011). As *sin4Δ* leads to *HO* hyperactivation and enhanced coactivator binding, we next wanted to investigate whether disrupting *SIN4* could also rescue *swi2* and *gcn5* coactivator mutants. To test this possibility, we examined Swi2-V5 and Gcn5-V5 binding when *SIN4* and either *GCN5* or *SWI2* were mutated, respectively.

As expected, disrupting *GCN5* alone resulted in a small decrease in Swi2-V5 at URS1, while disrupting *SIN4* alone resulted in a 3-fold increase in Swi2-V5 binding (Fig 7A). Disrupting *SIN4* and *GCN5* together yielded an ~3-fold increase in Swi2-V5 enrichment similar to that of the *sin4* single mutant. This result suggests that SWI/SNF binding at the *HO* promoter is no longer dependent on functional SAGA when *SIN4* is also disrupted. As a control, Swi2-V5 binding was similarly measured when *GAL11* was disrupted instead of *GCN5* and binding was found to be Tail-dependent (Fig 7A). We next examined Gcn5-V5 binding in a *swi2-314* (E843K) mutant that encodes a partially functional Swi2 protein (Mitra *et al*. 2006). Mutating *swi2* resulted in a large decrease in Gcn5-V5 enrichment, while disrupting *SIN4* resulted in a 50% increase in Gcn5-V5 binding over WT (Fig 7B). Combining *swi2-314* with *sin4Δ* rescued Gcn5-V5 binding to levels 20% higher than WT and nearly 2-fold higher than that found in the *swi2-314* strain alone. Taken together, these results indicate that recruitment of SWI/SNF and SAGA to the *HO* promoter are not required for each other when the Tail subcomplex is separated from the rest of Mediator.

**Fig 7.**
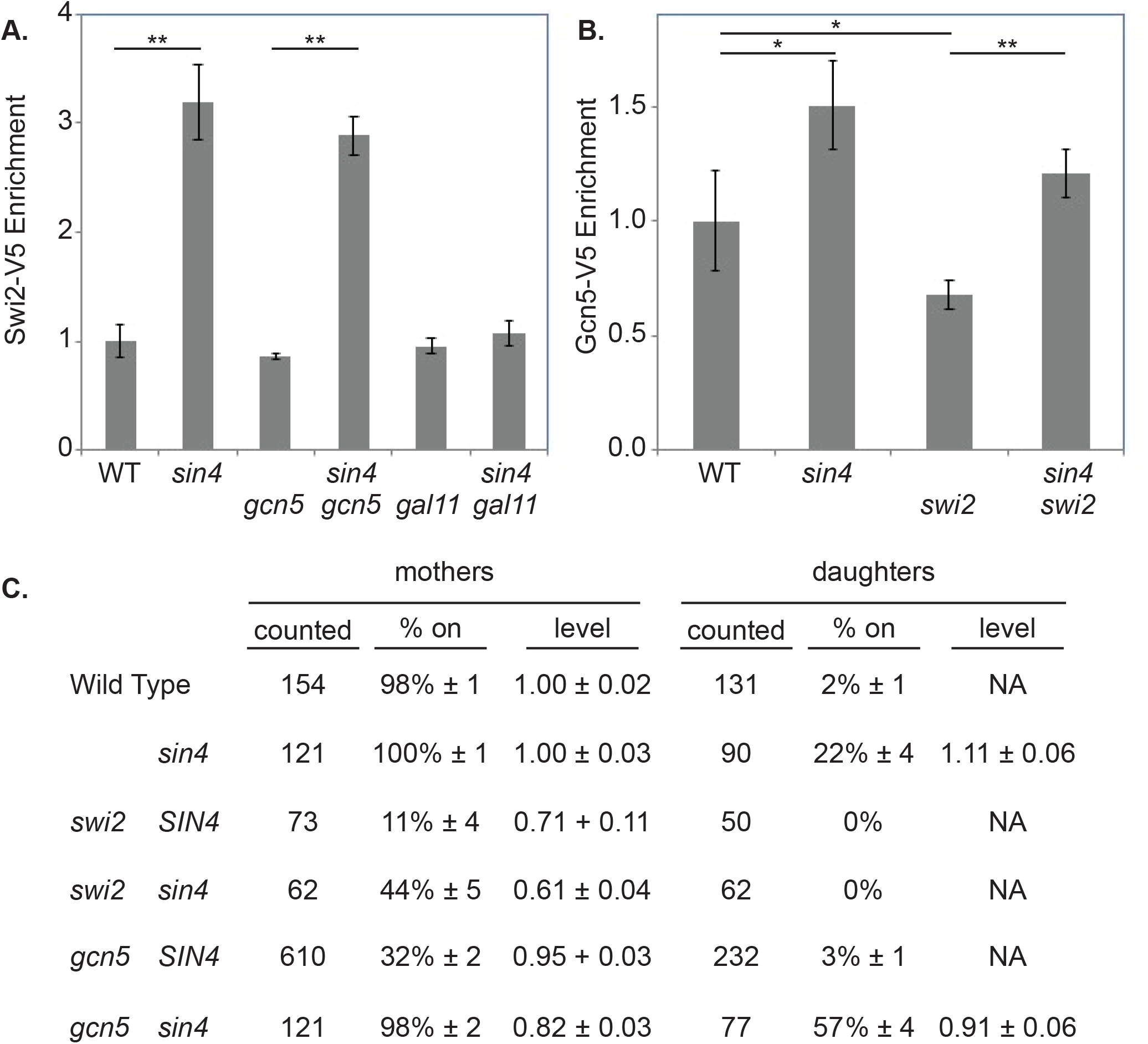
A *sin4* mutation suppresses defects at *HO* caused by coactivator mutations. **A.** A *sin4* mutation enhances SWI/SNF binding to *HO* despite a *gcn5* mutation. SWI/SNF binding to *HO* was measured by ChIP assays with Swi2-V5 in wild type, *sin4, gcn5, sin4 gcn5, gal11*, and *sin4 gal11* strains. The error bars reflect the standard deviation of three biological samples. \*\**p* < 0.01. **B.** A *sin4* mutation enhances the Gcn5-V5 binding to *HO* despite a *swi2-314* mutation. Gcn5 binding to *HO* was measured by ChIP assays in wild type, *sin4*, *swi2-314*, and *sin4 swi2-314* strains. The error bars reflect the standard deviation of three biological samples. \**p* < 0.05, \*\**p* < 0.01. **C.** A *sin4* mutation enhances *HO* expression despite coactivator mutations. Expression of an *HO-GFP* reporter was measured by single cell time-lapse fluorescence microscopy, in both mother and daughter cells. The table lists the number of cells counted, the percentage of cells in which *HO-GFP* was expressed, and the relative level of expression. Expression levels were normalized so that the average expression level in wild type mother cells is 1. The data for the wild type and *gcn5* mutants are from (Zhang *et al*. 2013b).

The defect in *HO* activation observed in *gcn5* and *swi2-314* cells could be due to WT or near-WT levels of activity from a subset of cells or to low levels of promoter activity from a larger fraction of cells. Examining these two options and how *sin4Δ* suppresses these mutants at the single cell level could provide additional information on the mechanism of its suppression. To address these possibilities, we combined *gcn5* and *swi2-314* mutations with an *HO-GFP* reporter and analyzed expression using single cell time-lapse fluorescence microscopy (Zhang et al., 2013). Single cell analysis revealed a significant reduction in the number of mother cells expressing at *HO-GFP* WT levels in both the *swi2* and *gcn5* single mutants, and this fraction increased significantly when a *sin4* mutation was introduced (Fig 7C). Disrupting *SIN4* did not result in cells expressing *HO* at levels surpassing WT, but rather increased the percentage of cells expressing *HO*. Independent recruitment of the Tail subcomplex therefore increases the probability of promoter activation within a population of cells. *HO* is normally expressed only in mother cells, and not in daughters (Jensen *et al*. 1983; Nasmyth 1983). Interestingly, the *sin4Δ gcn5Δ* double mutant displayed expression in a significant fraction of daughter cells, a property not found in either of the single mutants. Mechanistically, it is not at all clear why *HO* is expressed in the *sin4*Δ *gcn5*Δ double mutant, and this bears further investigation.

### Disruption of *SIN4* enables SWI/SNF binding under arrest conditions

A possible explanation for the loss of coactivator interdependence in *sin4Δ* is that the entire Mediator complex has both positive and negative roles in the binding of other coactivators. A similar argument has been proposed for SAGA, which is capable of stimulating SWI/SNF binding to chromatin via histone acetylation, but also facilitating SWI/SNF dissociation by direct Snf2 acetylation (Kim *et al*. 2010). By this model, the inhibitory roles of Mediator are limited to the core Mediator and the Kinase module, and are therefore absent when an independent Tail module alone is recruited to the promoter, thereby creating a permissive environment for other coactivators to bind. Elevated and persistent coactivator binding observed in *sin4Δ* (Fig 5C) supports this model.

To further explore this proposed model of inhibition by the complete Mediator complex, we next examined *HO* under arrest conditions in which only SBF and Mediator are bound to the promoter (Zhang *et al*. 2013b; Yu *et al*. 2016). If our model is correct, the lack of SWI/SNF binding under arrest conditions is at least partly due to inhibitory pressure from the kinase module and core Mediator complex that would be absent in *sin4*Δ. To test this theory, we examined Swi2 enrichment by Swi2-V5 ChIP under α-factor arrest in WT and *sin4Δ* cells. Since α-factor arrest induces expression of a lncRNA that disrupts factor binding (Yu *et al*. 2016), the cells used in this experiment have an *HO* promoter mutation at the Ste12 binding site required for inducing this lncRNA. Interestingly, while WT cells had Swi2-V5 levels comparable to no tag controls, *sin4Δ* had elevated Swi2-V5 binding to URS2L with 2.5-fold enrichment over background (Fig 8A). Although it is possible that this persistent Swi2 binding is due to the 3 to 4-fold greater binding that was observed during logarithmic growth, we saw nearly identical levels of Gal11 binding in WT and *sin4Δ* cells when we probed similarly for Gal11-V5 instead of Swi2-V5 (Fig 8B). These results support a model in which Mediator can both positively and negatively regulate SWI/SNF binding to promoters, and, furthermore, that these two regulatory effects can be separated from one another by independent recruitment of only the Tail module to promoters.

**Fig 8.**
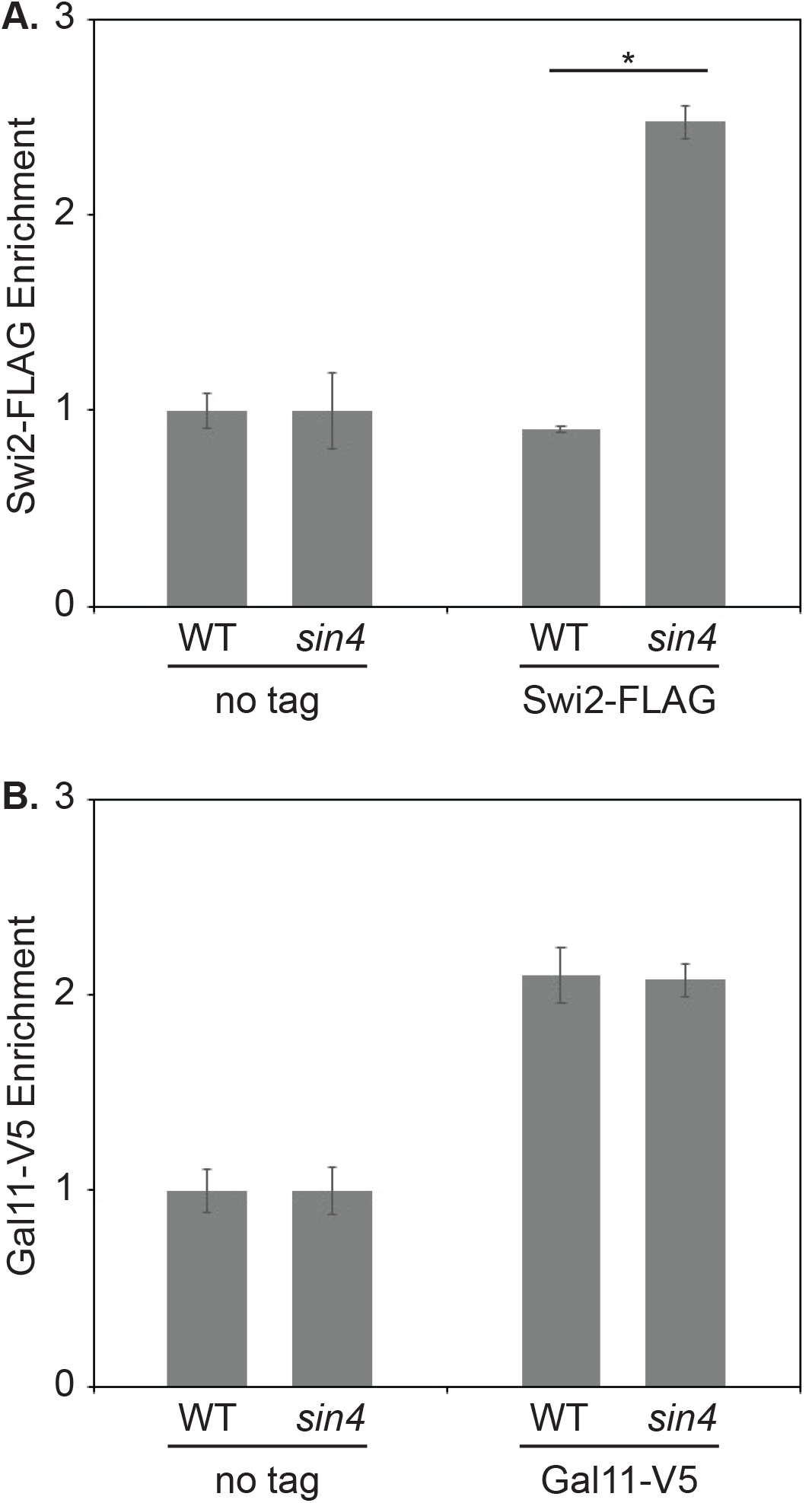
A *sin4* mutation facilitates SWI/SNF binding at *HO* during G1 arrest. **A.** SWI/SNF is normally not present at *HO* URS2 during a G1 arrest (Zhang *et al*. 2013b). A ChIP experiment shows that a *sin4* mutation allows Swi2-FLAG to bind to *HO* URS2 during a G1 arrest. The error bars reflect the standard deviation of two biological samples. \**p* < 0.05. **B.** A ChIP experiment shows that a *sin4* mutation does not result in an increase in Gal11-V5 binding to *HO* URS2 during a G1 arrest. The error bars reflect the standard deviation of two biological samples.

## DISCUSSION

Mediator consists of four conserved modules (Head, Middle, Kinase and Tail), and previous work has implicated the Kinase module as the primary antagonist to Tail binding at the UAS elements of genes (Van De Peppel *et al*. 2005; Jeronimo and Robert 2017). In this study, we probed the antagonist model at the *Saccharomyces cerevisiae HO* gene by utilizing a *SIN4* disruption that separates the Tail module from the Kinase module and core Mediator. In *sin4Δ* mutant cells we observed a significant increase in Tail module recruitment, elevated SWI/SNF and SAGA coactivator binding, persistent nucleosome eviction, and hyperactivation of the *HO* gene. These results are consistent with the Kinase module as the primary Tail antagonist, as the Kinase module would not be present at the UAS in the *sin4Δ* mutant to limit Tail module recruitment to the *HO* promoter. However, mutation of the catalytic subunit of the Kinase module failed to reproduce the hyperactivation of *HO* observed here with independent Tail recruitment. These results suggest that the activity and binding of the Tail is limited by other factors. One intriguing possibility is the core Mediator, which is also not present at the UAS during independent Tail module recruitment in the *sin4*Δ mutant, and thus core Mediator might function to limit Tail binding.

Our results suggest that the core Mediator may also limit the binding of the SWI/SNF and SAGA coactivators. At the *HO* promoter, coactivator binding is typically interdependent (Takahata *et al*. 2011), and impairment or loss of any one coactivator leads to diminished recruitment of the other coactivators and failure to express *HO* at significant levels. When the Tail module was recruited independently of the rest of Mediator, we observed greater than WT levels of SWI/SNF and SAGA coactivators, even when the other coactivator or its activity was disrupted. Additionally, when core Mediator was absent, we were able to observe considerable enrichment of SWI/SNF at *HO* URS2 even under α-factor arrest conditions in which coactivators are typically evicted by transcribing RNA polymerase (Yu *et al*. 2016). These results suggest that Mediator has both positive and negative roles in the recruitment of other coactivators, similar to those of SWI/SNF in the recruitment of the SAGA complex (Kim *et al*. 2010). We propose that core Mediator limits the ability of other coactivators to bind while the Tail module facilitates their binding.

An alternative explanation for the increased coactivator recruitment in *sin4*Δ is that the Tail subcomplex stimulates the recruitment of other coactivators, while core Mediator, along with the Kinase module, simply limits Tail occupancy at promoters. When *SIN4* was disrupted, we saw an ~50% increase in the enrichment of the Gal11 Tail subunit to *HO*. As coactivator binding is typically interdependent, this increase in Tail occupancy could be responsible for stimulating elevated recruitment of the SWI/SNF and SAGA coactivator complexes and rescuing recruitment of either of these coactivators when the other is impaired or disrupted. In this model, core Mediator would regulate coactivator binding by limiting the availability of the Tail module to the UAS or enhancers of genes.

Why would Mediator limit its own recruitment and the binding of other coactivators? The *Saccharomyces cerevisiae* genome is very compact with inter-ORF distances ranging from 150 to 400 bp (Pelechano *et al*. 2006). The increased coactivator recruitment and nucleosome eviction that we observe at *HO* when the Tail module is recruited independently has profound effects on *HO* transcriptional regulation, and it is possible that these effects are occurring genome wide. Although microarray analysis of *sin4Δ* does not reveal drastic changes in global transcript levels (Van De Peppel *et al*. 2005), changes in global chromatin structure and hypersensitivity to micrococcal nuclease have been reported (Macatee *et al*. 1997). We have shown previously that nucleosomes act as gates to regulate both activation and timing of expression (Yarrington *et al*. 2016), and coactivator recruitment and nucleosome eviction must be precise to limit promoter activation and ensure proper regulation of gene transcription.

Disruption of *SIN4* has been previously implicated in the long-range activation of genes (Dobi and Winston 2007). In this report, the authors identified *sin4Δ* as capable of enabling transcriptional activation at normally non-permissive distances of 800 bp or greater in *S. cerevisiae*. Interestingly, this effect was specific to *sin4*Δ as the authors were unable to reproduce long-range activation with other Mediator mutations affecting all four modules. These results are therefore consistent with the effect being due to an independent Tail subcomplex. Although the authors were unable to provide a mechanism for this altered activation by *sin4*Δ, based on our results with *HO*, it is likely that the observed long-range activation is due to independent Tail recruitment with associated elevated and persistent transcription factor and coactivator binding.

Lastly, it is important to note that we find no evidence that disrupting *SIN4* has altered the function of Mediator, only the recruitment of its four modules. Binding of the Tail subcomplex to *HO* still requires both the Gal11 Tail subunit and the Swi5 transcription factor, and *HO* expression requires SBF bound to URS2R. Furthermore, single cell analysis demonstrates that independent recruitment of the Tail subcomplex has not altered the mechanism of *HO* activation but rather the probability of activation. These results are consistent with whole Mediator working in concert with and regulating the recruitment of other coactivators for proper gene regulation.

## ACKNOWLEDGMENTS

We thank Tim Formosa and Emily Parnell for comments on the manuscript, and Tim Formosa and members of the Stillman lab for helpful advice throughout the course of this project. We thank Leena Bhoite, Sophie Song, and Warren Voth for providing strains with epitope tags, and Zaily Connell, Tim Formosa, John McCusker, and Elmar Schiebel for providing plasmids. This work was supported by National Institutes of Health grants GM121858 (L.B.) and GM121079 (D.J.S.)

---

**Supplemental Fig S1. The increased SBF binding in a *sin4* mutant persists during the cell cycle.**

Wild type (solid lines) and *sin4* (dotted lines) cells with a *GALp:CDC20* allele and a Swi4-V5 epitope tag were synchronized by galactose withdrawal and re-addition, and factor binding was measured by ChIP during the cell cycle. The ChIP signal is plotted as a function of time after release from the G2/M arrest. SBF binding was measured to the left (blue) or the right (red) parts of URS2, using the primers indicated on the diagram.

**Supplemental Fig S2. *sin4* and *ash1* mutations additively increase *HO* expression.**

*HO* mRNA levels were measured for the various mutant *HO* promoters indicated on the left, in either wild type or *sin4* mutants. The error bars reflect the standard deviation of two biological samples.

**Supplemental Fig S3. The Srb4 subunit of the Head module of Mediator is not recruited to *HO* in a *sin4* mutant.**

Srb4-V5 binding to *HO* URS1 and URS2 was measured by ChIP in wild type and a *sin4* mutant. The error bars reflect the standard deviation of three biological samples. \**p* < 0.05, \*\**p* < 0.01.

**Supplemental Fig S4. The *sin4* suppression of *HO* expression is dependent on the *PGD1* and *MED2* tail subunits.**

*HO* mRNA levels were measured in the indicated strains. The error bars reflect the standard deviation of two biological samples.

